# Brief Trains of Closed-loop TMS Induce Controllable Power in FM-theta Oscillations

**DOI:** 10.64898/2026.10.03.756435

**Authors:** Brian Erickson, Brian Kim, Alan Degenhart, Matti Stenroos, Ryan Rich, John Medaglia

## Abstract

**Objective:** Frontal-midline theta (FM-theta; 4–8 Hz) oscillations are thought to support cognition. Brief trains of transcranial magnetic stimulation (TMS) can probe these oscillations with high temporal precision but have rarely changed theta power. One possible reason is that prior studies ignored the instantaneous phase of the oscillation, which strongly shapes the cortical response to TMS. We tested whether brief TMS trains triggered at specific FM-theta phases in real time directionally modulate post-train FM-theta power.

**Approach:** Six healthy adults received TMS over dorsomedial prefrontal cortex (Fz) with concurrent electroencephalography during a whole-report working memory task. A beamformer and real-time phase-estimation pipeline triggered four-pulse 6 Hz trains (110% of resting motor threshold) at FM-theta peaks (0°) or troughs (180°), randomized trial by trial. Post-train spectral power was decomposed into total, induced (non-phase-locked) and evoked (phase-locked) components.

**Main results:** Trough-triggered trains transiently increased post-train FM-theta power relative to peak-triggered trains; task performance did not differ. Cluster-based permutation tests showed trough−peak differences in total (4.6–7.1 Hz) and induced (5.0–7.8 Hz) power ∼50–440 ms after the train, suggesting amplification of the oscillation . No evoked effect was detected. The total-power effect was robust at the subject level (Cohen’s dz = 1.32, p = 0.023). Within-train theta power did not differ between conditions, suggesting a cortical origin.

**Significance:** To our knowledge, this is the first study to lock brief rhythmic TMS trains to the real-time phase of an ongoing oscillation, and to show phase-dependent modulation of FM-theta power after individual trains on timescales relevant to behavior. Open-loop brief-train protocols may have missed such effects by mixing facilitatory and suppressive phases. Phase-locked brief trains offer a precise tool for testing the causal role of oscillations in cognition.

## Introduction

Frontal-midline theta (FM-theta; 4–8 Hz) power is associated with cognitive performance in health and disease across multiple domains, including working memory (WM), attention, and cognitive control (Cavanagh & Frank, 2014; Hsieh & Ranganath, 2014; Jensen & Tesche, 2002). FM-theta has accordingly been proposed as a neural mechanism supporting generalized executive control, potentially reflecting coordinated fronto-parietal network activity (Cavanagh & Frank, 2014; Sauseng et al., 2010). If FM-theta plays a causal role in cognition, then externally modulating this rhythm could reveal neurophysiological mechanisms and lead to therapies (Hanslmayr et al., 2019; Luber & Lisanby, 2014). Such interventions might be especially effective if they are sensitive to the real-time state of FM-theta and can optimally boost its power (Beynel et al., 2021).

Some neuromodulation strategies have been effective in influencing FM-theta. Most notably, repetitive transcranial magnetic stimulation (rTMS) can influence this rhythm and related behaviors (Hoy et al., 2016; Vékony et al., 2018), but these effects have been highly variable and often difficult to interpret (Beynel, Appelbaum, et al., 2019). One likely reason is a temporal mismatch between the stimulation and oscillatory process: conventional rTMS protocols deliver hundreds of pulses over many seconds and produce prolonged effects, whereas FM-theta itself is transient, task-dependent, and engaged by multiple distinct cognitive demands (Cavanagh & Frank, 2014).

To better match stimulation to the timescale of FM-theta, several studies have delivered brief trains of TMS - as few as four pulses to several dozen - at specific moments during cognitive tasks (Beynel, Davis, et al., 2019; Beynel et al., 2021; Luber et al., 2007, 2008; Riddle et al., 2020, 2024). These protocols have produced several significant effects on WM and related functions, establishing that brief trains can engage cognitively relevant processes. How they do so at the level of the targeted oscillation, however, remains unclear. The few brief-train studies with concurrent EEG have reported effects on phase locking, connectivity, and phase-amplitude coupling (Albouy et al., 2017; Glim et al., 2019; Riddle et al., 2024), but sustained power increases outlasting the train have rarely been observed. The only clear report used 18 pulses and targeted beta (Hanslmayr et al., 2014), while shorter trains have produced effects that dissipate quickly (Thut et al., 2011). The difficulty of inducing power enhancements is a significant limitation for brief-train studies, as many of the observed links between oscillations and cognition are power effects.

Beyond the number of pulses delivered, one reason brief-train protocols may have produced weak and inconsistent power effects in FM-theta and other oscillations is that they have not accounted for the instantaneous phase of the targeted oscillation. A growing body of evidence shows that TMS effects depend strongly on pre-pulse phase (Bergmann et al., 2019; Stefanou et al., 2018; Zrenner et al., 2018), because phase reflects ongoing fluctuations in cortical excitability. Phase effects can be large enough to produce a sign reversal, in which an intervention that enhances targeted oscillation at one phase suppresses it at the opposite phase (Zrenner et al., 2018). Accordingly, stimulation targeted to specific phases (often called “closed-loop” stimulation; cf. Bergmann, 2018; Karabanov et al., 2016) could have more powerful effects. Phase-targeted stimulation has been applied to FM-theta in only one prior study, which used rTMS rather than brief-trains (Gordon et al., 2022). To our knowledge, no study has used phase-targeted brief-train TMS in any frequency band. It therefore remains unknown whether locking brief trains of TMS to real-time oscillatory phase can produce reliable changes in post-train power at the trial level, in FM-theta or in other oscillations. If phase is a major source of this variability, phase-locked trains might produce reliable power effects with far fewer pulses than the 18-pulse train previously required. If it were possible to do so, this approach could unlock new experimental designs that test the causal roles of oscillations at the precise moments they are thought to be recruited to support cognition.

Accordingly, we delivered four-pulse 6 Hz TMS trains locked to either the peak or trough of ongoing FM-theta to test whether these conditions differentially modulate post-train theta power. Stimulation was embedded in a Whole Report (WR) working memory task (Adam et al., 2018). The phase condition randomized across WR trials. Because surface-EEG troughs (negative peaks) mark high-excitability states, at which TMS evokes larger corticospinal responses (Zrenner et al., 2018) and trough-locked prefrontal rTMS increases TMS-induced theta power (Gordon et al., 2022), stimulation at FM-theta troughs should enhance the endogenous oscillation. We hypothesized that trough-targeted stimulation would improve WM accuracy and transiently increase FM-theta power relative to peak-targeted stimulation in the immediate post-train interval. To test whether TMS modulated ongoing oscillatory amplitude independently of changes in cross-trial phase consistency, theta power was quantified separately for induced (non-phase-locked) and evoked (phase-locked) components.

## Methods

### Participants

We recruited 10 healthy adults (5 female; mean age = 31.7 years, SD = 12.8). All participants provided written informed consent. The study was approved by the Drexel University Institutional Review Board (protocol #1904007140) and conducted in accordance with the Declaration of Helsinki. All participants were screened for TMS contraindications (Rossi et al., 2021). Handedness was assessed using the Edinburgh Handedness Inventory (Oldfield, 1971; mean score = 88.8, SD = 21.6), and all included participants were right-handed.

Participants first completed a resting-state EEG screening session to determine eligibility for real-time phase targeting. Participants were required to exhibit a theta-band (4–8 Hz) spectral peak at electrode Fz of at least 2 dB above the fitted 1/f curve (via methods similar to Zrenner et al., 2020). Only those meeting this criterion proceeded to MRI scanning. Four individuals were excluded due to the absence of a clear frontal-midline theta (4–8 Hz) spectral peak. The final sample included 6 participants (2 female; mean age = 35.7 years, SD = 14.3, range 19–60; mean handedness score = 91.7, SD = 15.1). Sessions lasted approximately 2.5 hours, and participants were compensated $25/hour.

### Screening and Subject Preparation

Subjects participated in an initial screening EEG session, an MRI scan, and a TMS-EEG session. During the screening session, 20 minutes of resting-state EEG were recorded using a 64-channel actiCAP slim system with TMS-compatible Ag/AgCl active electrodes arranged according to the extended 10–10 system. Impedances were kept below 15 kΩ throughout the experiment. Electrode positions were digitized using Brainsight neuronavigation. EEG data were acquired in a shielded room via Lab Streaming Layer and the LSL-actiCHamp interface (Kothe et al., 2025). Resting-state data were analyzed offline to identify an individual frontal-midline theta (4–8 Hz) spectral peak. A linear constrained minimum variance (LCMV) beamformer was computed using head models generated from the MRI data and the 20 minutes of screener rest data (Gordon et al., 2021; Stenroos et al., 2007; Stenroos & Nummenmaa, 2016; Van Veen et al., 1997). Source orientation at each dmPFC target was fixed to the outward-pointing cortical surface normal. Each spatial filter was scaled to unit gain for its target’s lead field with the same sign convention in every participant.

### Phase Targeting Methods

EEG was streamed into a custom Python-based real-time pipeline (developed with Starfish Neuroscience) implementing an autoregressive (AR) prediction algorithm to estimate the future phase of ongoing FM-theta and trigger TMS at predefined phase targets (0° peak; 180° trough). AR prediction compensated for processing and hardware delays associated with EEG acquisition and TMS triggering (Figure 1A). A mean hardware lag of 12 ms was corrected online during targeting, with residual variability of SD = 2 ms. Beamformer output from the dmPFC source was used as the target channel for real-time phase estimation to improve signal-to-noise relative to channel-level EEG (Figure 1B).

**Figure 1:**
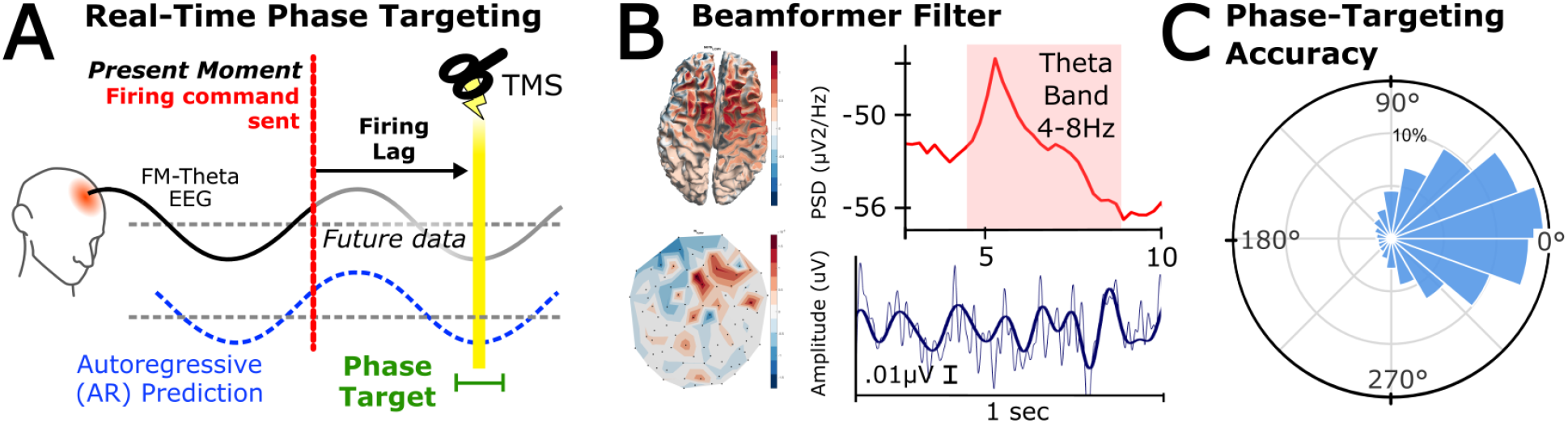
Real-Time FM-theta Phase Targeting and Validation. (A) Schematic of the phase-targeting framework. Frontal midline (FM) theta activity is acquired and processed in real time. The phase of the signal cannot be reliably estimated at the present moment due to filter boundary effects, and the processing loop time incurs lag. Accordingly, we used autoregressive prediction to estimate the future phase of the signal and issued the firing command in advance so that the pulse arrived at the intended phase target (here, trough). (B) Beamformer-based extraction of FM-theta. Left: spatial filter weights shown as scalp topography and corresponding cortical sensitivity profile. Right: example power spectral density demonstrating a theta-band peak (4–8 Hz), and representative time series illustrating broadband (1–55 Hz; thin light-blue line) and theta-band (4–8 Hz; thick dark-blue line) activity after spatial filtering. (C) Phase-targeting accuracy during simulation. Polar histogram shows the distribution of phase error (degrees) between the intended phase target (0°) and the realized phase at simulated pulse onset. The mean phase error (x̅), angular deviation (σ), and number of events (N) are indicated. Concentration of events around 0° demonstrates accurate targeting of the desired FM-theta phase.

TMS artifacts prevent direct estimation of instantaneous phase at the time of stimulation (Zrenner et al., 2018). Accordingly, targeting accuracy was evaluated using simulated firing events (Figure 1C). The real-time pipeline was re-applied to the rest data offline using that subject’s dmPFC beamformer, the session’s live power thresholds and a 100ms post-trigger timeout. Triggers were simulated at both peak and trough phase targets. At each simulated trigger the real-time estimate was compared with a ground-truth phase computed offline by filtering the entire recording (4–8 Hz acausal zero-phase Butterworth filter, Hilbert transform). Group-level accuracy, defined as the proportion of triggers falling within ±45° of the ground-truth phase, was 55.0%. Group level circular mean error and the angular deviation √(2(1−R)) was 10.6° (±55.4°). This phase-targeting precision is comparable to prior FM-theta phase-targeted implementations (Gordon et al., 2021); Supplementary Figure 1).

### TMS-EEG Methods

After beamformer computation, participants completed a TMS-EEG session. Stimulation was delivered over Fz (10–20 system), corresponding to dmPFC (Gordon et al., 2021), while participants performed a discrete whole-report (WR) working memory task.

Participants were semi-reclined, fitted with a 96-channel EEG cap and earplugs, and registered to their individual MRI-derived head model using neuronavigation. Resting motor threshold (RMT) was determined using a neuronavigated Magstim D70 coil. RMT was defined as the minimum intensity eliciting motor-evoked potentials ≥50 μV in the left first dorsal interosseous muscle on 5 of 10 pulses (Schutter & van Honk, 2006). Stimulation intensity was set to 110% of RMT to induce reliable cortical effects (Rossini et al., 2015).

Participants first completed a 4-minute eyes-open resting-state block (used for phase-targeting validation), followed by the stimulation block. The WR task followed (Adam et al., 2018), with the addition of phase-targeted TMS synchronized to stimulus presentation (Figure 2). The task was implemented in PsychoPy (Peirce et al., 2019). On each trial, a green arrow cue indicated the relevant hemifield, followed by bilateral arrays of six colored squares per hemifield presented for 250 ms. After a 1,200 ms retention interval, participants reported the color at each of the six cued locations by selecting from a color grid; responses were unspeeded, and trials were separated by a 1 s inter-trial interval. Participants completed three to five runs of approximately 60 trials each.

**Figure 2:**
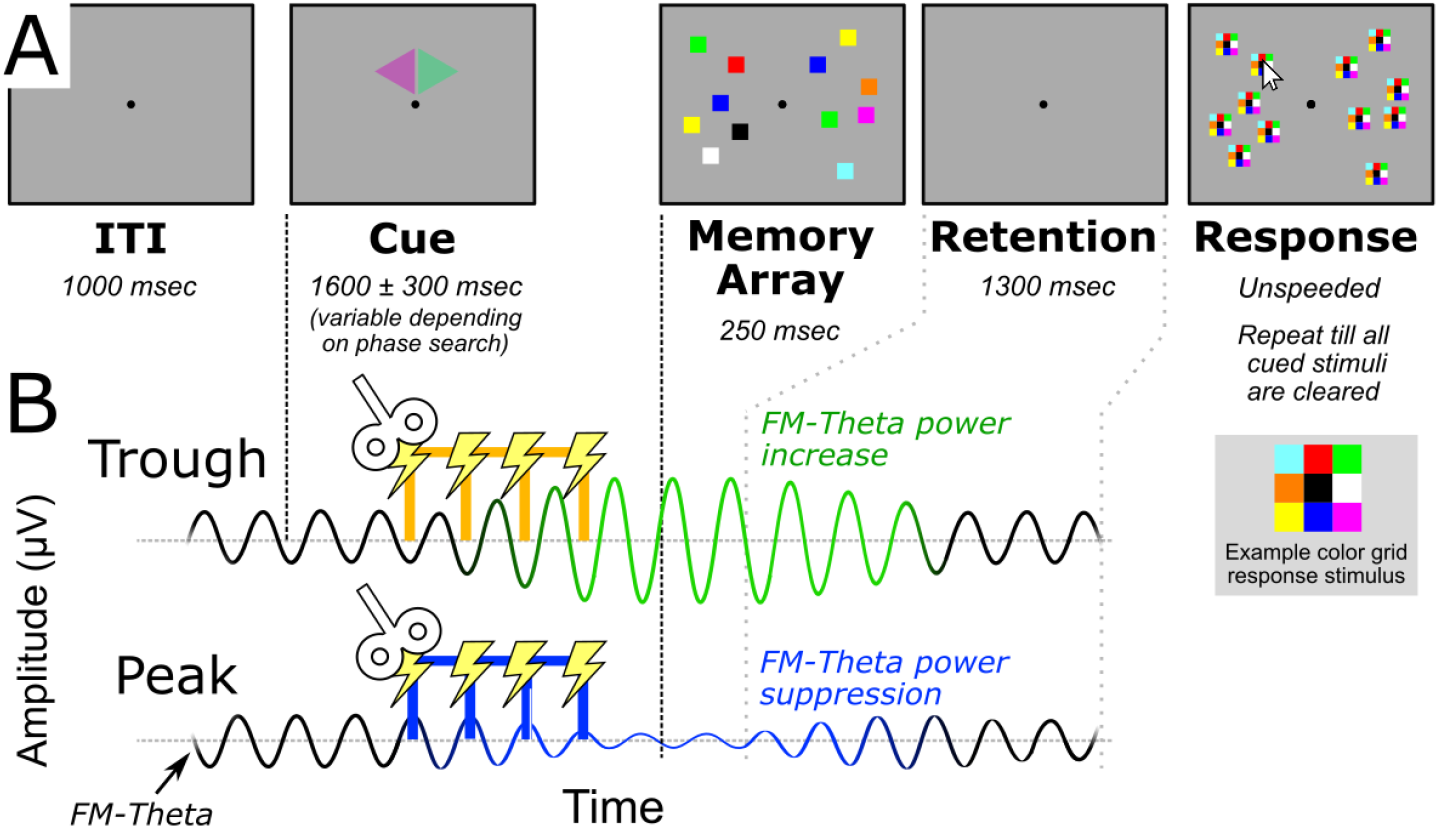
Whole-Report Task and Phase-Targeted Stimulation Design. (A) Discrete whole-report (WR) working memory task. A spatial cue (green arrow) indicated the relevant hemifield, followed by presentation of bilateral arrays of six colored squares. After a delay, participants reported remembered colors by selecting from a color grid at each location (stimuli enlarged for clarity). Responses were unspeeded. (B) Phase-targeted stimulation design. During the cue period, the real-time pipeline triggered a four-pulse 6 Hz TMS train at either the peak (0°) or trough (180°) of ongoing FM-theta, contingent on detection of a valid phase target. The memory array was presented 700ms following train offset. We hypothesized that trough-targeted stimulation would produce a transient post-train increase in FM-theta power relative to peak-targeted stimulation.

During the cue period preceding the memory array at 700ms, PsychoPy sent a network command enabling the phase-targeting pipeline to trigger stimulation at the next acceptable peak (0°) or trough (180°). Triggering additionally required that the mean theta-band (4–8 Hz) amplitude of the beamformer signal fall within a session-specific window set by the experimenter, whose upper bound excluded high-amplitude periods likely to reflect artifacts. If a valid phase target was detected within the next 600 ms, a firing command was issued and PsychoPy paused 500 ms to allow completion of the four-pulse 6 Hz train. If no valid target was detected, the trial proceeded without stimulation. Valid targets were detected on 80 to 90% of trials for all subjects. The cue remained on the screen for an additional 700ms before presentation of the memory array stimuli.

Phase condition (peak vs trough) was randomized on a trial-by-trial basis, and both participant and experimenter were blind to condition. TMS was positioned over Fz using a mechanical arm and monitored throughout the session (Magstim D70 Air Film, Superrapid 2+1 unit, equivalent output to D70 coil used to determine RMT).

### TMS-EEG Data Preprocessing and Analysis

EEG preprocessing focused on minimizing TMS-related artifacts to permit analysis of post-train theta power. Bad channels and noisy segments were first rejected from the continuous data. Pulse artifacts were processed using the TESA toolbox for EEGLAB (Delorme & Makeig, 2004; Rogasch et al., 2017). For each pulse, data within ±10 ms of pulse onset were removed, cubically interpolated using a ±20 ms window surrounding the cut segment, bandpass filtered (1–50 Hz), and the ±10 ms pulse window then set to zero again so that the interpolated placeholder used for filtering was not retained as data. Data were then re-referenced to the common average of all retained channels, previously rejected channels interpolated (spherical spline), and independent component analysis (ICA) was performed. Components identified as artifactual by ICLabel (Pion-Tonachini et al., 2019; default settings) and manual inspection were removed. Cleaned channel-level data were spatially filtered using the subject-specific beamformer to extract the dmPFC source signal. Data were epoched from −1000 to 2000 ms relative to the final pulse in each train to capture the pre-pulse baseline, pulse train and a sufficient period to evaluate the full evolution of post-train spectral power. Trial metadata (peak vs trough target) were coded as EEG flags. Then, time-frequency analysis was performed separately for peak- and trough-targeted trials using EEGLAB’s event-related spectral perturbation (ERSP) function. Power was decomposed into three components, computed separately for each condition. Total power was obtained by applying the wavelet transform to each single trial and averaging power across trials. Evoked (phase-locked) power was obtained by applying the same transform to the condition’s across-trial average waveform. Induced (non-phase-locked) power was obtained by subtracting the condition’s average waveform from every trial before applying the transform and averaging power across trials. ERSPs were baselined from −400 to −10 ms prior to the start of the pulse train, to capture a sufficient period of power for a stable baseline while avoiding any filter artifacts remaining after TESA pulse interpolation. Wavelet decomposition began with 3 cycles at 3 Hz, increasing linearly to 5.6 cycles at 14 Hz (EEGLAB newtimef, cycles = [3 0.6]), with single-trial gain model normalization (Grandchamp & Delorme, 2011). Trough−peak differences were assessed with a cluster-based permutation test on subject-level difference maps (trough minus peak) within an a priori window of 4–8 Hz and 550–1200 ms (50–700 ms after train offset), separately for total, induced and evoked power. At each time-frequency bin, a paired t statistic was computed across subjects; bins exceeding the cluster-defining threshold (|t(5)| > 2.571, two-tailed p < .05) were grouped into clusters by 8-neighbour adjacency, and cluster mass was defined as the summed t value. The null distribution of the maximum absolute cluster mass was obtained from all 2⁶ = 64 possible sign inversions of the subjects’ difference maps (an exact test; equivalent to exchanging condition labels within subjects), and clusters were considered significant at a family-wise α of 0.05. The same test was applied to theta power during the pulse train (0–500 ms).

## Results

Phase-targeted stimulation differentially modulated frontal-midline theta (FM-theta) power. The cluster-based permutation test over the a priori window (4–8 Hz, 550–1200 ms; see Methods) revealed significant clusters for both total and induced power. The total power cluster spanned 4.6–7.1 Hz beginning 53 ms post-train offset with a duration of 390 ms (53–443 ms post-offset; cluster mass = 1053.96, p = 0.031). The induced power cluster spanned 5.0–7.8 Hz over the same time extent (cluster mass = 1170.45, p = 0.031; Figure 3). Both of these p values equal 2/64, the smallest attainable with six participants in our sign-flip permutation test. The analysis window began at 50 ms post-train offset to exclude residual TMS pulse artifacts; thus, the significant clusters begin near the start of the analysis window. No significant clusters emerged for evoked power, nor for total theta band power over the pulse train interval (0 to 500 ms; smallest cluster p = 0.19), indicating that within-train theta increases did not differ between conditions. Subject-level and average ERSPs plotted over an expanded frequency window (3–14 Hz) are included in Supplementary Figure 2.

**Figure 3:**
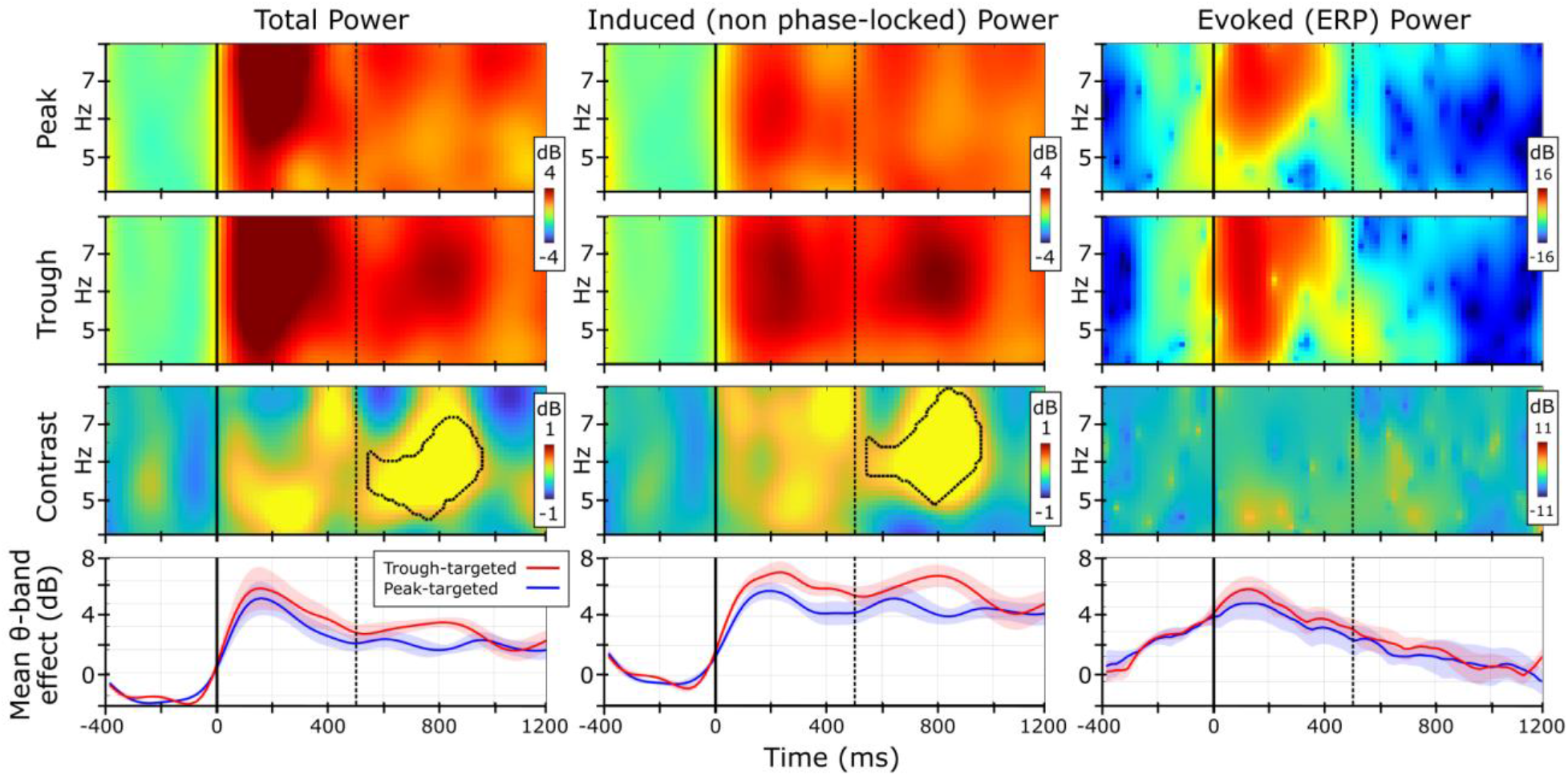
Group Average Event-Related Spectral Perturbations (ERSPs). Top row: peak-targeted stimulation. Second row: trough-targeted stimulation. Third row: trough–peak contrast, masked by cluster-based permutation significance (exact sign-flip permutation test, familywise error corrected at p < 0.05). Bottom row: mean theta-band ERSP across frequency bins; shaded regions denote standard error of the mean. Columns (left to right): total power, induced (non-phase-locked) power, and evoked (phase-locked) power. The TMS train began at zero (solid vertical black line) and ended at 500ms (dashed vertical line). A significant increase in total and induced theta power for trough compared to peak stimulation was observed beginning immediately after train offset (at the 50ms post-train artifact cut window) and lasting approximately 400ms.

Total and induced power effects were most prominent in the middle of theta band (near 6 Hz), although the precise peak frequency varied across individuals (Figure 4). To complement the cluster-based group analysis with a subject-level effect size, we extracted each participant’s mean dB difference (trough − peak) within the same a-priori window as the cluster permutation test. Total power yielded a robust effect (mean = 0.49 dB, SD = 0.37, t(5) = 3.25, p = 0.023, Cohen’s dz = 1.32, 95% t-based CI on mean difference [0.10, 0.87] dB) indicating that our stimulation effect generalizes across participants and converging with our cluster permutation test. We did not detect evoked power effects (mean = 0.88 dB, SD = 3.77, 95% CI [−3.07, 4.84] dB; t(5) = 0.57, p = 0.59, dz = 0.23). Induced power showed a similar mean difference as total power (0.54 dB, SD = 0.57, 95% CI [−0.06, 1.14] dB) but was trending at the subject level (t(5) = 2.31, p = 0.069, dz = 0.94). This result in induced power likely underestimates the true effect for two reasons. First, our long a-priori defined time-frequency region of interest (ROI) included approximately 300ms after the total and induced power effects dissipated (as defined by the cluster test) which biased our results towards the null. In a descriptive post-hoc window spanning the approximate extent of the significant group-level clusters (4–8 Hz, 550–900 ms), the subject-level induced effect was significant (mean = 0.81 dB, SD = 0.56, 95% CI [0.22, 1.40] dB; t(5) = 3.51, p = 0.017, dz = 1.43; Figure 4, bottom right panel), as was the total power effect (mean = 0.79 dB, SD = 0.69, 95% CI [0.07, 1.52] dB; t(5) = 2.80, p = 0.038, dz = 1.15), whereas evoked power again showed no difference (t(5) = 0.73, p = 0.50). Second, the two subjects with the weakest overall theta-band dB effect in the a priori ROI (S2 and S6) both showed isolated strong negative contrast effects in the a priori ROI. These effects were centered in adjacent bands — Delta for S2 and Alpha for S6 — but overlapped the theta band (Figure 4). This spectral leakage attenuated the overall theta-band effect for these subjects. Leakage is greatly reduced for S2 in the post-hoc ROI, as it occurred late in the epoch.

**Figure 4:**
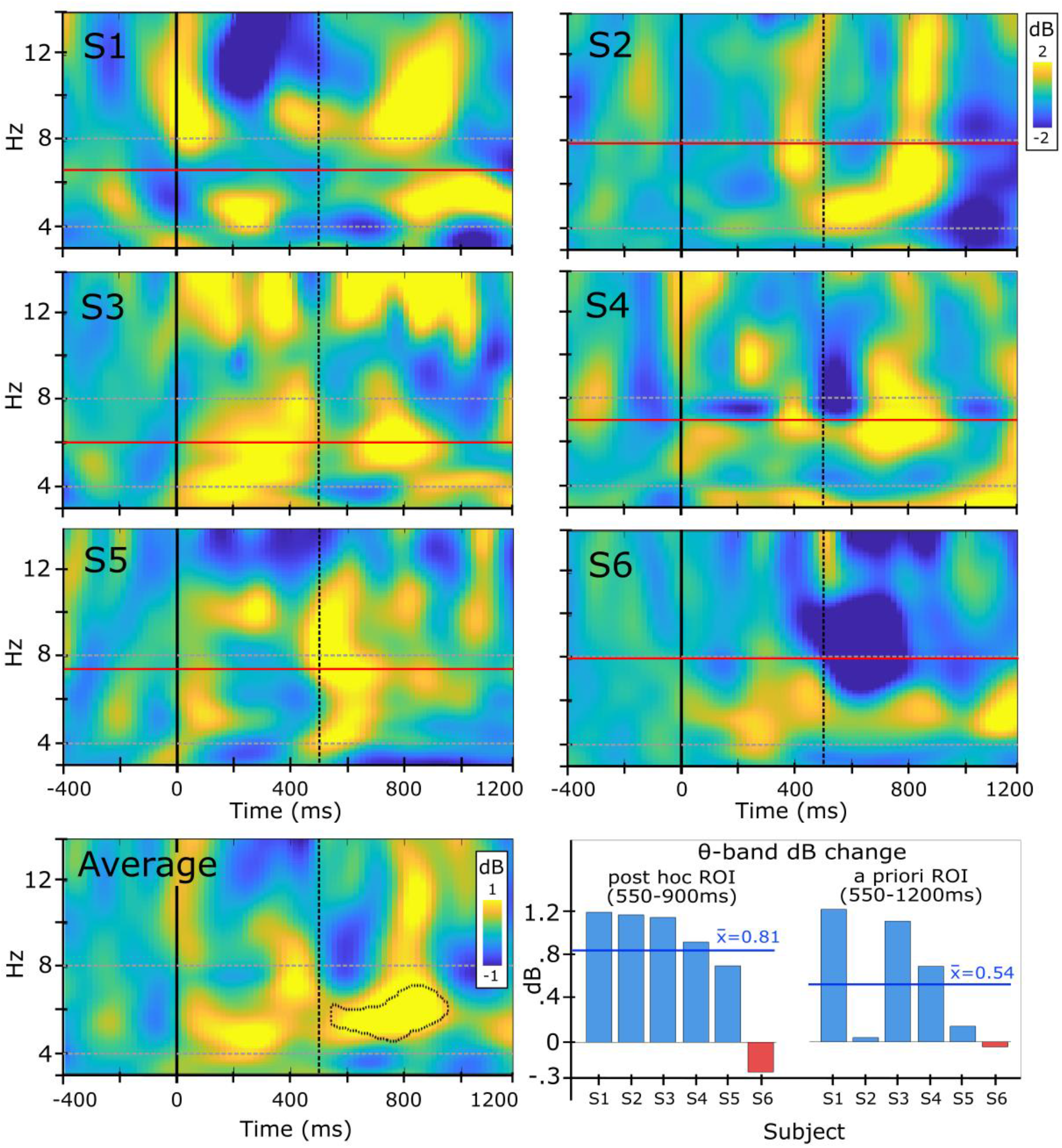
Subject-Level Induced Power Contrast (Trough − Peak). *Upper six panels:* induced event-related spectral perturbation (ERSP) contrast power is shown for each individual subject (S1–S6) plotted over an expanded frequency range. Power is expressed as baseline-normalized change (dB). The TMS train began at 0 ms (solid vertical black line) and ended at 500 ms (dashed vertical line). Horizontal dashed lines denote theta band (4–8 Hz). Horizontal solid red line denotes each subject’s peak theta frequency calculated from the rest data used to compute the beamformer. *Bottom left panel:* group-average induced contrast map, with the significant cluster from the a priori window (4–8 Hz, 550–1200 ms) outlined. *Bottom right panel:* induced theta-band dB contrast effect per subject averaged over time-frequency regions of interest defined post hoc (left) and a priori (right).

Despite the effect on FM-theta power, TMS phase targeting condition did not predict accuracy or reaction time on the WR task. Notably, our design presented the WM stimuli approximately 250ms beyond the end of the brief-train theta power effect. We modeled behavioral outcomes with mixed-effects models (Phase: Peak vs. Trough; random intercept for Subject) and compared subject-level measures (Cowan’s K, median RT) with paired t-tests. Accuracy, defined as the proportion of correctly identified items among the six probed locations per trial, did not differ by phase (Peak: 43.8%; Trough: 43.1%; binomial GLMM, β = −0.04, SE = 0.05, t(1156) = −0.78, p = .438), nor did Cowan’s K (K = N(pc − pg)/(1 − pg); Peak: M = 2.21; Trough: M = 2.16; t(5) = −0.72, p = .502). Response time to the first item on each trial likewise did not differ by phase (Peak: M = 1.49 s; Trough: M = 1.43 s; t(1154) = −1.29, p = .197). Subject-level median RT trended faster on Trough versus Peak trials (1.26 vs. 1.35 s, a ∼92 ms / 7% difference; t(5) = −2.20, p = .079, dz = 0.90); this large effect did not reach significance with the present sample, possibly due to the limited N (6 subjects).

## Discussion

Brief trains of TMS (four pulses) enhanced post-train theta power when targeted to troughs relative to peaks of the endogenous frontal-midline theta (FM-theta) rhythm. The enhancement was present in induced (non-phase-locked) power, indicating that the stimulation amplified the ongoing oscillation. Our findings support that endogenous phase influences the neural effects of brief rhythmic TMS trains, and motivate that future causal studies using such TMS trains should consider and control the influence of phase on their stimulation. To our knowledge, our study is the first to lock brief rhythmic TMS trains to the real-time phase of an ongoing oscillation during task trials, and the first demonstration that brief trains of TMS produce transient phase-dependent modulation of FM-theta power.

Brief trains of rhythmic TMS have been shown to affect cognition (Albouy et al., 2017; Chanes et al., 2013; Hanslmayr et al., 2014; Riddle et al., 2020; Romei et al., 2012) and measures such as EEG connectivity (Stengel et al., 2021), phase locking (Albouy et al., 2017; Vernet et al., 2019), phase-amplitude coupling (Glim et al., 2019), and frontoparietal theta connectivity (Riddle et al., 2024). However, reports of post-train power increases in this literature have been rare, weak, or depended on a high pulse count (18 pulses; Hanslmayr et al., 2014). This is surprising, as the most basic expectation after a high energy rhythmic influence like TMS might be that the targeted oscillation would become bigger. Our results suggest that the absence of such power effects in prior studies may be because those studies were “open-loop”, that is, did not control for FM-theta phase. If endogenous phase influences how TMS affects oscillatory power, as our study suggests, open-loop designs inherently mix trials at facilitatory and ineffective or even suppressive phases. This would bias those studies towards null power effects. This interpretation aligns with a seminal study by Thut et al., 2011, which showed that brief-train power depends on post-hoc phase. Our study extends this result to FM-theta and demonstrates that prospectively conditioning stimulation “closed-loop” on endogenous phase influences effects on post-train power. The induced power effect was highly reliable (5/6 subjects) when accounting for our overly long a-priori ROI (Figure 4, bottom right panel).

Our results agree with the popular notion that EEG phase reflects cyclic fluctuations in cortical excitability. Stimulation at troughs could coincide with periods of heightened depolarization, thereby amplifying ongoing activity, whereas peak stimulation may coincide with relatively refractory states. Such state-dependence was originally demonstrated for motor cortex alpha oscillations (Bergmann et al., 2019; Stefanou et al., 2018; Zrenner et al., 2018) and later extended to FM-theta oscillations and WM in a closed-loop rTMS design (Gordon et al., 2022). Our study used similar methodology to target FM-theta phase (Gordon et al., 2021), but applied brief trains of stimulation. The punctate (∼400ms) post-train power effect we demonstrated could be used to manipulate FM-theta power in real-time, on individual trials, to test how FM-theta oscillations causally support cognitive function. Specifically, studies could time this brief modulation of power to occur at the moment an oscillation is theorized to support cognition. A possible reason we did not observe effects on WM behavior in the present study is that we presented the WM array approximately 250ms after the theta power effect dissipated. Future experiments can use the timing of the power effect we identified as a guide for experimental design.

We found an increase in induced FM-theta power, which is evidence that our stimulation directly affected the power of the endogenous FM-theta oscillation. Oscillatory power can appear to change in response to TMS for two reasons. First, TMS can straightforwardly make the amplitude of the oscillation bigger or smaller. However, a second reason is that TMS can “reset” the endogenous oscillation to a consistent phase (Erickson et al., 2024). Then when the EEG is averaged over trials to create a “TMS evoked potential” (TEP), this TEP will be larger wherever the phase was more consistent, even if the amplitude of the underlying oscillation did not change (Makeig et al., 2002; Sauseng et al., 2007). These effects can accumulate even in open-loop designs, because TMS can reset oscillations to a consistent phase regardless of their starting point (Erickson et al., 2024). Notably, the majority of prior brief-train studies have exclusively detected evoked, phase-related effects that could be driven by resets (Albouy et al., 2017; Glim et al., 2019; Riddle et al., 2024). Induced power is not subject to this ambiguity because it subtracts the ERP from each trial before power computation. Thus, the induced power effect we found likely reflects a genuine amplification of the endogenous oscillation: aligning phase across trials without changing amplitude would shift power from the induced to the evoked component, and so could not increase induced power. Whether stimulation also altered cross-trial phase alignment remains open, because evoked power estimates were highly variable across participants, and phase-locked responses to a fixed 6 Hz train would also differ with each participant’s peak theta frequency.

With further study, it could be possible to finely tune the effects of rhythmic stimulation. Dynamic systems theory provides tools to understand how stimulation parameters influence the effects on endogenous oscillations. In particular, rhythmic stimulation only entrains (or “captures”) endogenous phase when the input is within a certain regime of duration, amplitude, and difference from the endogenous frequency (“detuning”) known as the “Arnold tongue” (Glass, 2001; Herrmann et al., 2016; Thut et al., 2011). Our stimulation was short (four pulses), slightly above RMT (110%), and at a fixed frequency (6 Hz). This may have placed our stimulation parameters outside the Arnold tongue, and as a result, induced an “amplitude response” (power enhancement; Castejón & Guillamon, 2013) with no detectable phase resetting. Speculatively, but consistent with this notion, our participants with peak theta frequencies closer to 6 Hz showed stronger power entrainment (Figure 4). This observation parallels recent evidence that individual resting-state spectral properties predict susceptibility to rhythmic TMS entrainment (Trajkovic et al., 2024). Whether an amplitude response or phase entrainment effect is preferable is a matter of experimental design. For instance, it could be important to not reset endogenous phase to avoid disrupting interregional phase coupling, or, driving changes in such coupling could be a study’s intended intervention. Therefore, we would ideally map how different TMS input parameters influence endogenous oscillations to enable comprehensive control (Herrmann et al., 2016).

The phase-targeted conditions we applied are excellent internal controls for each other, and our results are thus unlikely to reflect peripheral nervous system artifacts, differences in alertness, or other confounds. Peak and trough-targeted trials were identical in every respect except the phase target, randomized at the trial level, and passed the same real-time trigger constraints. The state of endogenous FM-theta is unlikely to have influenced peripheral artifacts, which is supported by a lack of any difference in within-train theta power. Together, these controls and results suggest a cortical origin of effect.

The FM-theta power effect we observed was highly consistent across subjects. Our subject-level tests converged with our group-level cluster analysis and had a large effect size (dz = 1.32). Potentially, such an effect size could enable designs at low enough trial counts to study a variety of FM-theta-linked cognitive functions (Cavanagh & Frank, 2014; Hsieh & Ranganath, 2014; McLoughlin et al., 2022). Our pre-planned analysis ROI spanned a conservatively long duration after the train, which diluted the induced power result. The true subject-level induced effect may thus be larger and more reliable than our a priori test suggests, though confirmation will require an independent sample. Prior tests of FM-theta’s contribution to WM have relied on designs such as long trains, rTMS, and correlational EEG-behavior relationships that cannot fully isolate the causal contribution of oscillatory state from the many other factors governing a trial’s outcome. Our phase-targeted brief-train approach is, to our knowledge, the first tool with the temporal precision to put this question to a direct test.

### Limitations

While we focused on a small, well-controlled design with the goal to enhance and identify effects within each individual, replication in a larger cohort will be important to establish the robustness and generalizability of the neurophysiological effects. In addition, four of the ten screened participants (40%) lacked a clear resting FM-theta peak and could not be phase-targeted, so our findings apply to individuals with a detectable FM-theta rhythm. Notably, the stimulation frequency was fixed at 6 Hz rather than individualized to each participant’s peak FM-theta frequency, which may have limited the magnitude or duration of entrainment in some individuals. Optimal performance may require tracking and stimulating at the instantaneous frequency of the endogenous oscillator, for instance, using state-space Kalman filtering approaches (Wodeyar et al., 2021). The study compared peak- and trough-targeted stimulation but did not include a sham or no-stimulation condition. Accordingly, the data establish a phase-dependent difference between the two stimulation conditions, but not versus any “baseline” condition. Although the trial-level randomization and matched stimulation parameters argue against a simple peripheral explanation, residual auditory, somatosensory, or muscle-related contributions cannot be completely excluded. Our study identified only the post-train enhancement window for FM-theta; other frequency bands and cortical generators may show different temporal dynamics that remain to be characterized. Future studies should align task events with the empirically identified modulation window to determine whether these transient oscillatory changes influence cognition.

## Conclusion

The instantaneous phase of an endogenous frontal-midline theta rhythm systematically conditions the short-lived neural response to a four-pulse rhythmic TMS train. Relative to peak-targeted stimulation, trough-targeted stimulation produced greater post-train theta power for several hundred milliseconds, with effects evident in total and induced power. This establishes endogenous phase as an experimentally useful control variable for brief rhythmic TMS and identifies a subsecond window in which phase-targeted stimulation can perturb oscillatory dynamics for future causal studies. In neurophysiological and cognitive studies, this approach offers excellent internal control because neural phase is inherently hidden from participants, effectively blinding them to the experimental condition on each trial. Furthermore, it is possible that this approach may generalize to other EEG oscillations. Accordingly, phase-locked stimulation may improve the specificity and reliability of oscillation-targeted neuromodulation, with potential implications for advancing control neural engineering in basic research and clinical applications. Whether this approach can reliably alter behavior, generalize across rhythms and cortical systems, or benefit from individualized frequency targeting remains to be tested.

## Acknowledgements

Funding: This work was supported by the National Institutes of Health [to J D M; grant number DP5-OD-021352]; and philanthropic donations from Starfish Neuroscience, Inc., Bellevue, WA [to J D M].

The authors acknowledge the use of Claude Opus 4.5–5.5 (Anthropic) to support improvements in internal consistency and compliance with journal guidelines, and for checking and refining analysis scripts. All AI-suggested text and code was critically revised by the authors. The final research remains our own responsibility.

## Abbreviations

AR: Autoregressive
dmPFC: Dorsomedial Prefrontal Cortex
EEG: Electroencephalogram
ERSP: Event Related Spectral Perturbation
LCMV: Linearly Constrained Minimum Variance
RMT: Resting Motor Threshold
TEPs: Transcranial-Evoked Potential components
TMS: Transcranial Magnetic Stimulation
WM: Working Memory
WR: Whole Report

## Supplementary Information

**Supplementary Figure 1:**
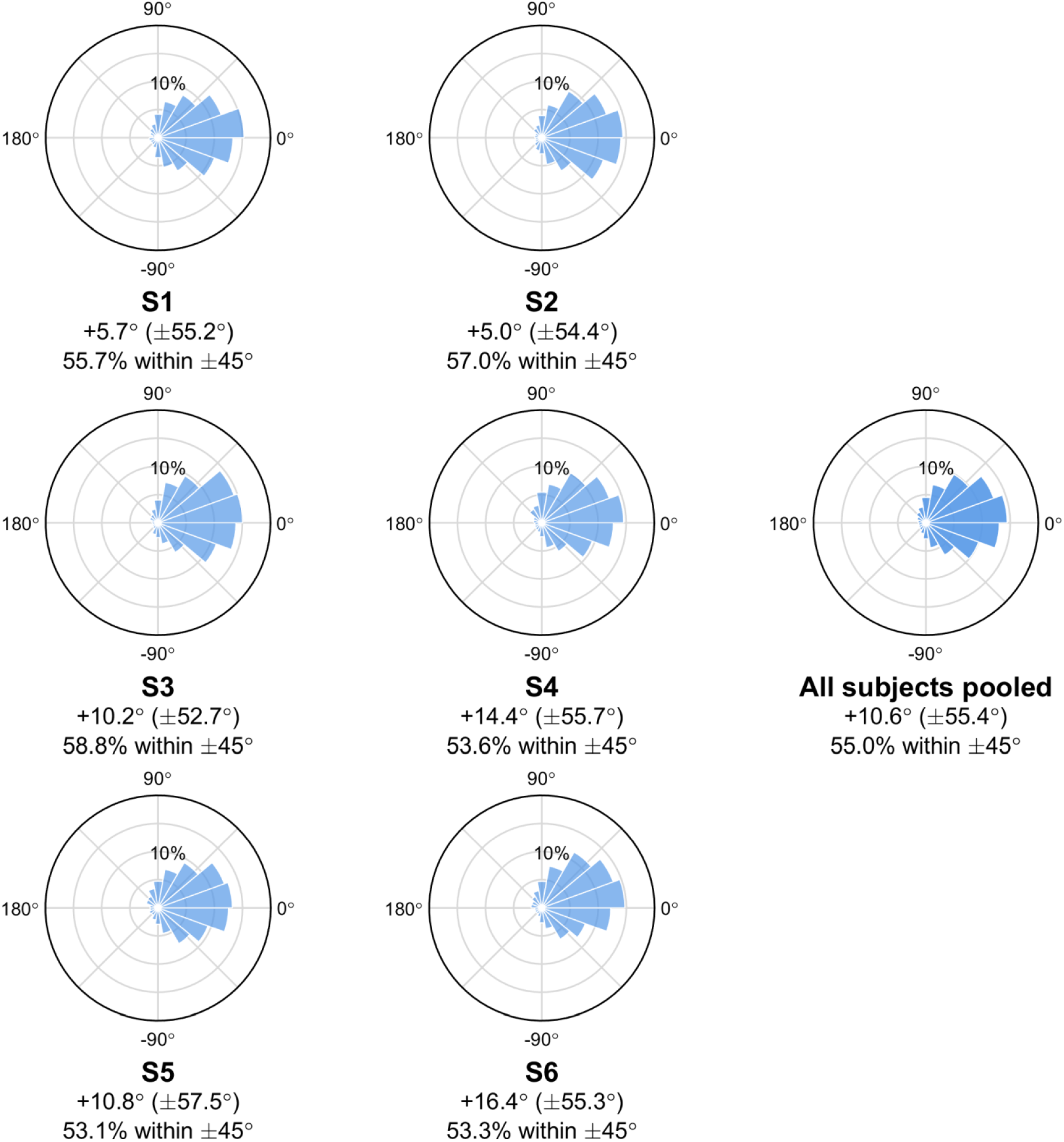
Individual Phase-Targeting Accuracy. Polar histograms of phase error (intended 0° target vs. simulated realized phase at pulse onset) for each subject (S1-S6), computed as described in the caption of Figure 1C. Below each plot is reported accuracy defined as the proportion of triggers falling within ±45° of the ground-truth phase, and circular mean error and the angular deviation defined as √(2(1−R)).

**Supplementary Figure 2:**
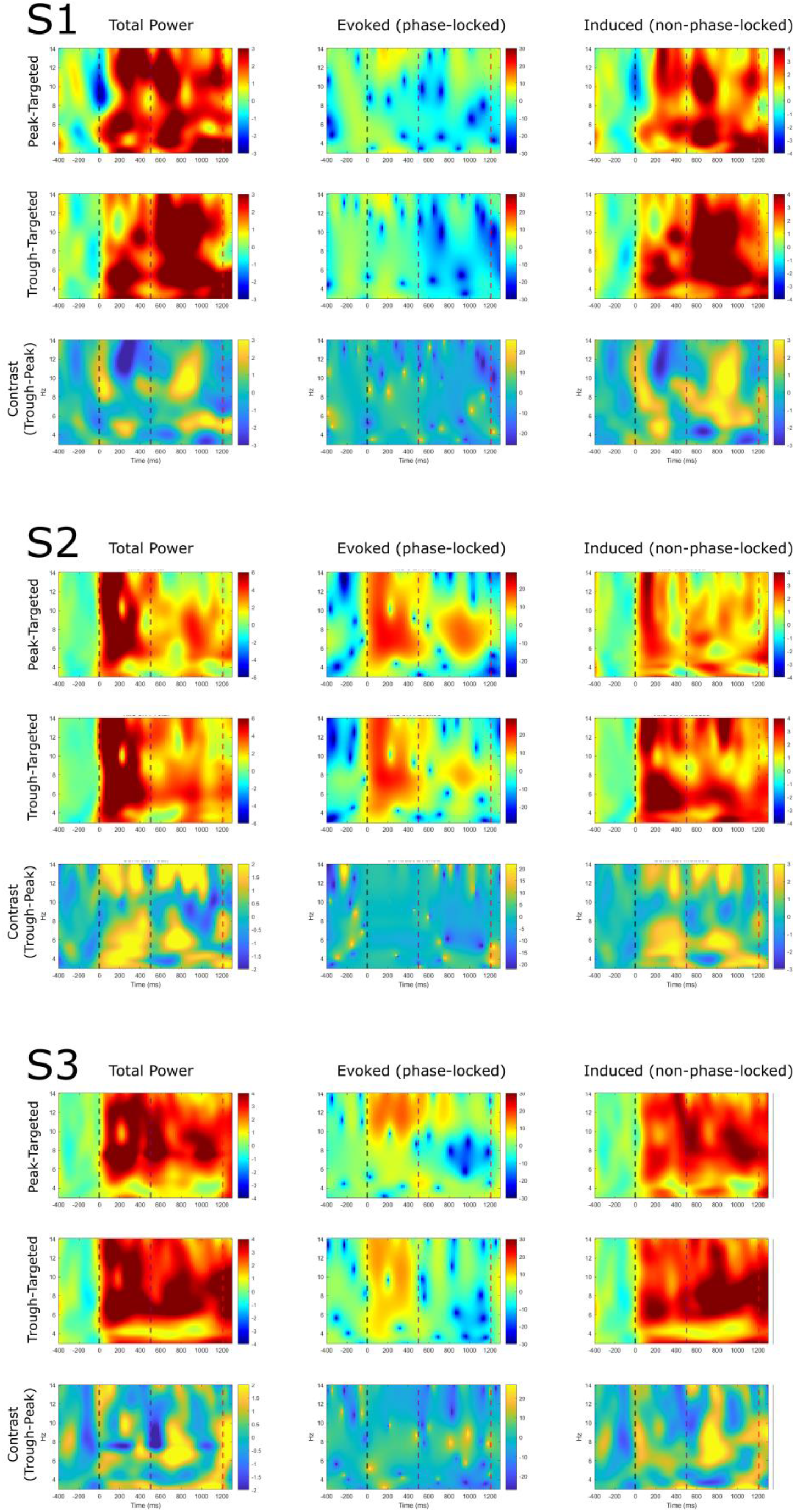
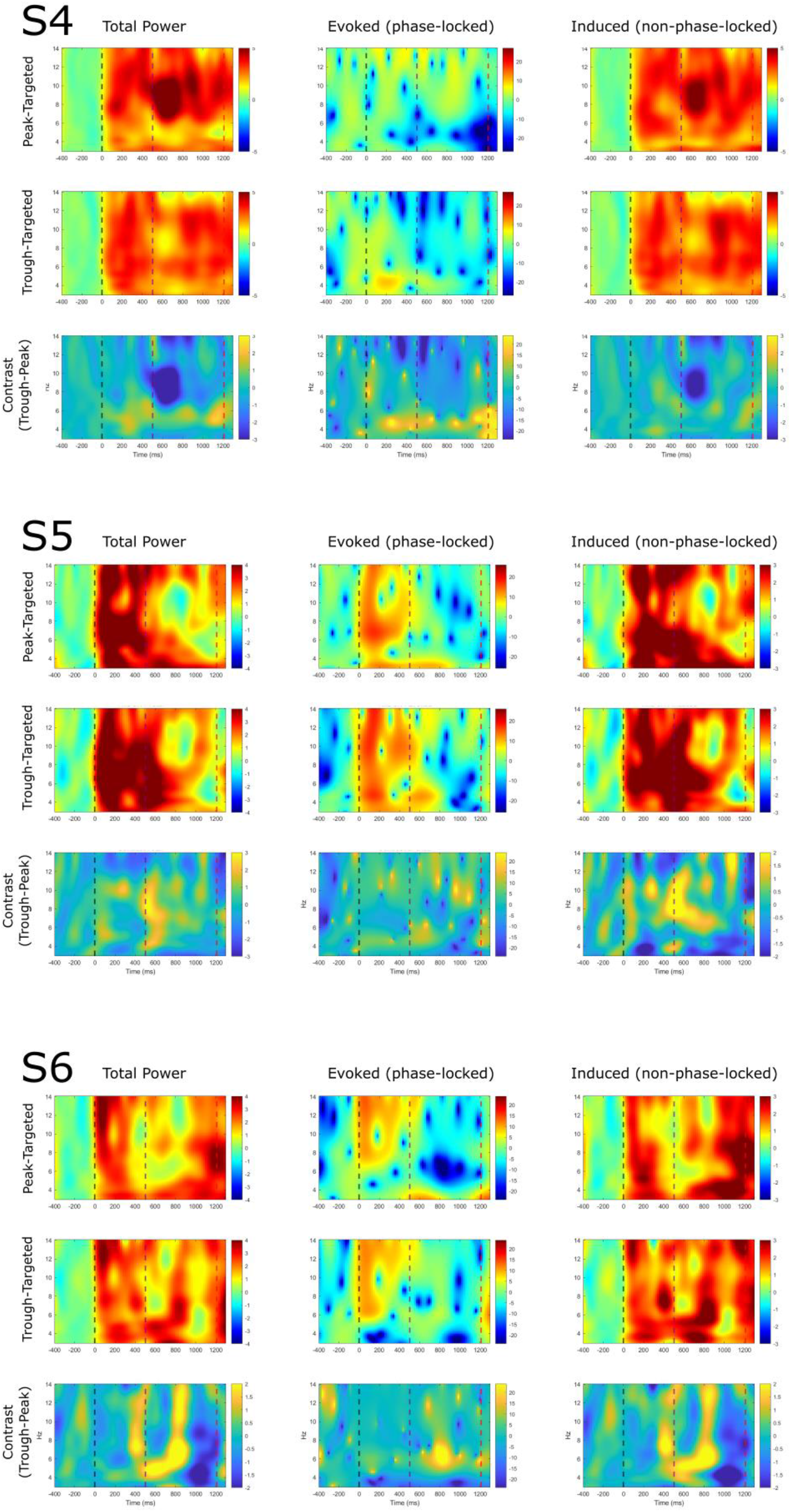
Subject-Level Time-Frequency Decomposition. ERSPs for each subject (S1–S6) over an expanded frequency range, decomposed into total power, evoked (phase-locked) power, and induced (non-phase-locked) power (columns), shown separately for peak-targeted trials, trough-targeted trials, and their contrast (trough-peak; rows). Vertical dashed lines mark train onset (0 ms) and offset (500 ms) and the onset of the memory stimulus (1200 ms)

## References

1. Adam, K. C. S., Robison, M. K., & Vogel, E. K. (2018). Contralateral Delay Activity Tracks Fluctuations in Working Memory Performance. Journal of Cognitive Neuroscience, 30(9), 1229–1240. 10.1162/jocn_a_01233

2. Albouy, P., Weiss, A., Baillet, S., & Zatorre, R. J. (2017). Selective Entrainment of Theta Oscillations in the Dorsal Stream Causally Enhances Auditory Working Memory Performance. Neuron, 94(1), 193–206.e5. 10.1016/j.neuron.2017.03.015

3. Bergmann, T. O. (2018). Brain state-dependent brain stimulation. Frontiers in Psychology, 9, 2108.

4. Bergmann, T. O., Lieb, A., Zrenner, C., & Ziemann, U. (2019). Pulsed Facilitation of Corticospinal Excitability by the Sensorimotor μ-Alpha Rhythm. The Journal of Neuroscience, 39(50), 10034–10043. 10.1523/JNEUROSCI.1730-19.2019

5. Beynel, L., Appelbaum, L. G., Luber, B., Crowell, C. A., Hilbig, S. A., Lim, W., Nguyen, D., Chrapliwy, N. A., Davis, S. W., Cabeza, R., Lisanby, S. H., & Deng, Z.-D. (2019). Effects of online repetitive transcranial magnetic stimulation (rTMS) on cognitive processing: A meta-analysis and recommendations for future studies. Neuroscience & Biobehavioral Reviews, 107, 47–58. 10.1016/j.neubiorev.2019.08.018

6. Beynel, L., Dannhauer, M., Palmer, H., Hilbig, S. A., Crowell, C. A., Wang, J. E.-H., Michael, A. M., Wood, E. A., Luber, B., Lisanby, S. H., Peterchev, A. V., Cabeza, R., Davis, S. W., & Appelbaum, L. G. (2021). Network-based rTMS to modulate working memory: The difficult choice of effective parameters for online interventions. Brain and Behavior, 11(11), e2361. 10.1002/brb3.2361

7. Beynel, L., Davis, S. W., Crowell, C. A., Hilbig, S. A., Lim, W., Nguyen, D., Palmer, H., Brito, A., Peterchev, A. V., Luber, B., Lisanby, S. H., Cabeza, R., & Appelbaum, L. G. (2019). Online repetitive transcranial magnetic stimulation during working memory in younger and older adults: A randomized within-subject comparison. PLOS ONE, 14(3), e0213707. 10.1371/journal.pone.0213707

8. Castejón, O., & Guillamon, A. (2013). Phase-Amplitude Response Functions for Transient-State Stimuli. Journal of Mathematical Neuroscience, 3, 13. 10.1186/2190-8567-3-13

9. Cavanagh, J. F., & Frank, M. J. (2014). Frontal theta as a mechanism for cognitive control. Trends in Cognitive Sciences, 18(8), 414–421. 10.1016/j.tics.2014.04.012

10. Chanes, L., Quentin, R., Tallon-Baudry, C., & Valero-Cabré, A. (2013). Causal Frequency-Specific Contributions of Frontal Spatiotemporal Patterns Induced by Non-Invasive Neurostimulation to Human Visual Performance. The Journal of Neuroscience, 33(11), 5000–5005. 10.1523/JNEUROSCI.4401-12.2013

11. Delorme, A., & Makeig, S. (2004). EEGLAB: An open source toolbox for analysis of single-trial EEG dynamics including independent component analysis. Journal of Neuroscience Methods, 134(1), 9–21. 10.1016/j.jneumeth.2003.10.009

12. Erickson, B., Kim, B., Sabes, P., Rich, R., Hatcher, A., Fernandez-Nuñez, G., Mentzelopoulos, G., Vitale, F., & Medaglia, J. (2024). TMS-induced phase resets depend on TMS intensity and EEG phase. Journal of Neural Engineering. 10.1088/1741-2552/ad7f87

13. Glass, L. (2001). Synchronization and rhythmic processes in physiology. Nature, 410(6825), 277–284. 10.1038/35065745

14. Glim, S., Okazaki, Y. O., Nakagawa, Y., Mizuno, Y., Hanakawa, T., & Kitajo, K. (2019). Phase-Amplitude Coupling of Neural Oscillations Can Be Effectively Probed with Concurrent TMS-EEG. Neural Plasticity, 2019, 1–13. 10.1155/2019/6263907

15. Gordon, P. C., Belardinelli, P., Stenroos, M., Ziemann, U., & Zrenner, C. (2022). Prefrontal theta phase-dependent rTMS-induced plasticity of cortical and behavioral responses in human cortex. Brain Stimulation, 15(2), 391–402. 10.1016/j.brs.2022.02.006

16. Gordon, P. C., Dörre, S., Belardinelli, P., Stenroos, M., Zrenner, B., Ziemann, U., & Zrenner, C. (2021). Prefrontal Theta-Phase Synchronized Brain Stimulation With Real-Time EEG-Triggered TMS. Frontiers in Human Neuroscience, 15, 691821. 10.3389/fnhum.2021.691821

17. Grandchamp, R., & Delorme, A. (2011). Single-Trial Normalization for Event-Related Spectral Decomposition Reduces Sensitivity to Noisy Trials. Frontiers in Psychology, 2. 10.3389/fpsyg.2011.00236

18. Hanslmayr, S., Axmacher, N., & Inman, C. S. (2019). Modulating Human Memory via Entrainment of Brain Oscillations. Trends in Neurosciences, 42(7), 485–499. 10.1016/j.tins.2019.04.004

19. Hanslmayr, S., Matuschek, J., & Fellner, M.-C. (2014). Entrainment of Prefrontal Beta Oscillations Induces an Endogenous Echo and Impairs Memory Formation. Current Biology, 24(8), 904–909. 10.1016/j.cub.2014.03.007

20. Herrmann, C. S., Murray, M. M., Ionta, S., Hutt, A., & Lefebvre, J. (2016). Shaping Intrinsic Neural Oscillations with Periodic Stimulation. The Journal of Neuroscience, 36(19), 5328–5337. 10.1523/JNEUROSCI.0236-16.2016

21. Hoy, K. E., Bailey, N., Michael, M., Fitzgibbon, B., Rogasch, N. C., Saeki, T., & Fitzgerald, P. B. (2016). Enhancement of Working Memory and Task-Related Oscillatory Activity Following Intermittent Theta Burst Stimulation in Healthy Controls. Cerebral Cortex, 26(12), 4563–4573. 10.1093/cercor/bhv193

22. Hsieh, L.-T., & Ranganath, C. (2014). Frontal midline theta oscillations during working memory maintenance and episodic encoding and retrieval. NeuroImage, 85, 721–729. 10.1016/j.neuroimage.2013.08.003

23. Jensen, O., & Tesche, C. D. (2002). Frontal theta activity in humans increases with memory load in a working memory task. European Journal of Neuroscience, 15(8), 1395–1399. 10.1046/j.1460-9568.2002.01975.x

24. Karabanov, A., Thielscher, A., & Siebner, H. R. (2016). Transcranial brain stimulation: Closing the loop between brain and stimulation. Current Opinion in Neurology, 29(4), 397–404.

25. Kothe, C., Shirazi, S. Y., Stenner, T., Medine, D., Boulay, C., Grivich, M. I., Artoni, F., Mullen, T., Delorme, A., & Makeig, S. (2025). The lab streaming layer for synchronized multimodal recording. Imaging Neuroscience, 3, IMAG.a.136. 10.1162/IMAG.a.136

26. Luber, B., Kinnunen, L. H., Rakitin, B. C., Ellsasser, R., Stern, Y., & Lisanby, S. H. (2007). Facilitation of performance in a working memory task with rTMS stimulation of the precuneus: Frequency- and time-dependent effects. Brain Research, 1128, 120–129. 10.1016/j.brainres.2006.10.011

27. Luber, B., & Lisanby, S. H. (2014). Enhancement of human cognitive performance using transcranial magnetic stimulation (TMS). NeuroImage, 85, 961–970. 10.1016/j.neuroimage.2013.06.007

28. Luber, B., Stanford, A. D., Bulow, P., Nguyen, T., Rakitin, B. C., Habeck, C., Basner, R., Stern, Y., & Lisanby, S. H. (2008). Remediation of Sleep-Deprivation-Induced Working Memory Impairment with fMRI-Guided Transcranial Magnetic Stimulation. Cerebral Cortex, 18(9), 2077–2085. 10.1093/cercor/bhm231

29. Makeig, S., Westerfield, M., Jung, T.-P., Enghoff, S., Townsend, J., Courchesne, E., & Sejnowski, T. J. (2002). Dynamic Brain Sources of Visual Evoked Responses. Science, 295(5555), 690–694. 10.1126/science.1066168

30. McLoughlin, G., Gyurkovics, M., Palmer, J., & Makeig, S. (2022). Midfrontal Theta Activity in Psychiatric Illness: An Index of Cognitive Vulnerabilities Across Disorders. Biological Psychiatry, Biomarkers of Psychosis, 91(2), 173–182. 10.1016/j.biopsych.2021.08.020

31. Oldfield, R. C. (1971). The assessment and analysis of handedness: The Edinburgh inventory. Neuropsychologia, 9(1), 97–113. 10.1016/0028-3932(71)90067-4

32. Peirce, J., Gray, J. R., Simpson, S., MacAskill, M., Höchenberger, R., Sogo, H., Kastman, E., & Lindeløv, J. K. (2019). PsychoPy2: Experiments in behavior made easy. Behavior Research Methods, 51(1), 195–203. 10.3758/s13428-018-01193-y

33. Pion-Tonachini, L., Kreutz-Delgado, K., & Makeig, S. (2019). ICLabel: An automated electroencephalographic independent component classifier, dataset, and website. NeuroImage, 198, 181–197. 10.1016/j.neuroimage.2019.05.026

34. Riddle, J., McPherson, T., Sheikh, A., Shin, H., Hadar, E., & Frohlich, F. (2024). Internal Representations Are Prioritized by Frontoparietal Theta Connectivity and Suppressed by alpha Oscillation Dynamics: Evidence from Concurrent Transcranial Magnetic Stimulation EEG and Invasive EEG. The Journal of Neuroscience, 44(15), e1381232024. 10.1523/JNEUROSCI.1381-23.2024

35. Riddle, J., Scimeca, J. M., Cellier, D., Dhanani, S., & D’Esposito, M. (2020). Causal Evidence for a Role of Theta and Alpha Oscillations in the Control of Working Memory. Current Biology, 30(9), 1748–1754.e4. 10.1016/j.cub.2020.02.065

36. Rogasch, N. C., Sullivan, C., Thomson, R. H., Rose, N. S., Bailey, N. W., Fitzgerald, P. B., Farzan, F., & Hernandez-Pavon, J. C. (2017). Analysing concurrent transcranial magnetic stimulation and electroencephalographic data: A review and introduction to the open-source TESA software. NeuroImage, 147, 934–951. 10.1016/j.neuroimage.2016.10.031

37. Romei, V., Thut, G., Mok, R. M., Schyns, P. G., & Driver, J. (2012). Causal implication by rhythmic transcranial magnetic stimulation of alpha frequency in feature-based local vs. global attention: Causal role of alpha in featural attention. European Journal of Neuroscience, 35(6), 968–974. 10.1111/j.1460-9568.2012.08020.x

38. Rossi, S., Antal, A., Bestmann, S., Bikson, M., Brewer, C., Brockmöller, J., Carpenter, L. L., Cincotta, M., Chen, R., Daskalakis, J. D., Di Lazzaro, V., Fox, M. D., George, M. S., Gilbert, D., Kimiskidis, V. K., Koch, G., Ilmoniemi, R. J., Lefaucheur, J. P., Leocani, L., … Hallett, M. (2021). Safety and recommendations for TMS use in healthy subjects and patient populations, with updates on training, ethical and regulatory issues: Expert Guidelines. Clinical Neurophysiology, 132(1), 269–306. 10.1016/j.clinph.2020.10.003

39. Rossini, P. M., Burke, D., Chen, R., Cohen, L. G., Daskalakis, Z., Di Iorio, R., Di Lazzaro, V., Ferreri, F., Fitzgerald, P. B., George, M. S., Hallett, M., Lefaucheur, J. P., Langguth, B., Matsumoto, H., Miniussi, C., Nitsche, M. A., Pascual-Leone, A., Paulus, W., Rossi, S., … Ziemann, U. (2015). Non-invasive electrical and magnetic stimulation of the brain, spinal cord, roots and peripheral nerves: Basic principles and procedures for routine clinical and research application. An updated report from an I.F.C.N. Committee. Clinical Neurophysiology, 126(6), 1071–1107. 10.1016/j.clinph.2015.02.001

40. Sauseng, P., Griesmayr, B., Freunberger, R., & Klimesch, W. (2010). Control mechanisms in working memory: A possible function of EEG theta oscillations. Neuroscience & Biobehavioral Reviews, 34(7), 1015–1022. 10.1016/j.neubiorev.2009.12.006

41. Sauseng, P., Klimesch, W., Gruber, W. R., Hanslmayr, S., Freunberger, R., & Doppelmayr, M. (2007). Are event-related potential components generated by phase resetting of brain oscillations? A critical discussion. Neuroscience, 146(4), 1435–1444. 10.1016/j.neuroscience.2007.03.014

42. Schutter, D. J. L. G., & van Honk, J. (2006). A Standardized Motor Threshold Estimation Procedure for Transcranial Magnetic Stimulation Research. The Journal of ECT, 22(3), 176. 10.1097/01.yct.0000235924.60364.27

43. Stefanou, M.-I., Desideri, D., Belardinelli, P., Zrenner, C., & Ziemann, U. (2018). Phase Synchronicity of μ-Rhythm Determines Efficacy of Interhemispheric Communication Between Human Motor Cortices. The Journal of Neuroscience, 38(49), 10525–10534. 10.1523/JNEUROSCI.1470-18.2018

44. Stengel, C., Vernet, M., Amengual, J. L., & Valero-Cabré, A. (2021). Causal modulation of right hemisphere fronto-parietal phase synchrony with Transcranial Magnetic Stimulation during a conscious visual detection task. Scientific Reports, 11(1), 3807. 10.1038/s41598-020-79812-y

45. Stenroos, M., Mäntynen, V., & Nenonen, J. (2007). A Matlab library for solving quasi-static volume conduction problems using the boundary element method. Computer Methods and Programs in Biomedicine, 88(3), 256–263. 10.1016/j.cmpb.2007.09.004

46. Stenroos, M., & Nummenmaa, A. (2016). Incorporating and Compensating Cerebrospinal Fluid in Surface-Based Forward Models of Magneto- and Electroencephalography. PLOS ONE, 11(7), e0159595. 10.1371/journal.pone.0159595

47. Thut, G., Veniero, D., Romei, V., Miniussi, C., Schyns, P., & Gross, J. (2011). Rhythmic TMS Causes Local Entrainment of Natural Oscillatory Signatures. Current Biology, 21(14), 1176–1185. 10.1016/j.cub.2011.05.049

48. Trajkovic, J., Sack, A. T., & Romei, V. (2024). EEG-based biomarkers predict individual differences in TMS-induced entrainment of intrinsic brain rhythms. Brain Stimulation, 17(2), 224–232. 10.1016/j.brs.2024.02.016

49. Van Veen, B. D., Van Drongelen, W., Yuchtman, M., & Suzuki, A. (1997). Localization of brain electrical activity via linearly constrained minimum variance spatial filtering. IEEE Transactions on Biomedical Engineering, 44(9), 867–880. 10.1109/10.623056

50. Vékony, T., Németh, V. L., Holczer, A., Kocsis, K., Kincses, Z. T., Vécsei, L., & Must, A. (2018). Continuous theta-burst stimulation over the dorsolateral prefrontal cortex inhibits improvement on a working memory task. Scientific Reports, 8(1), 14835. 10.1038/s41598-018-33187-3

51. Vernet, M., Stengel, C., Quentin, R., Amengual, J. L., & Valero-Cabré, A. (2019). Entrainment of local synchrony reveals a causal role for high-beta right frontal oscillations in human visual consciousness. Scientific Reports, 9(1), 14510. 10.1038/s41598-019-49673-1

52. Wodeyar, A., Schatza, M., Widge, A. S., Eden, U. T., & Kramer, M. A. (2021). A state space modeling approach to real-time phase estimation. eLife, 10, e68803. 10.7554/eLife.68803

53. Zrenner, C., Desideri, D., Belardinelli, P., & Ziemann, U. (2018). Real-time EEG-defined excitability states determine efficacy of TMS-induced plasticity in human motor cortex. Brain Stimulation, 11(2), 374–389. 10.1016/j.brs.2017.11.016

54. Zrenner, C., Galevska, D., Nieminen, J. O., Baur, D., Stefanou, M.-I., & Ziemann, U. (2020). The shaky ground truth of real-time phase estimation. NeuroImage, 214, 116761. 10.1016/j.neuroimage.2020.116761

